# An evolutionarily conserved Hox-Gbx segmentation code in the rice coral *Montipora capitata*

**DOI:** 10.1101/2024.09.29.615694

**Authors:** Shuonan He, Emma Rangel-Huerta, Eric Hill, Lacey Ellington, Shiyuan (Cynthia) Chen, Sofia Robb, Eva Majerová, Crawford Drury, Matthew C. Gibson

**Affiliations:** Stowers Institute for Medical Research, Kansas City, Missouri 64110, USA; Hawaiʻi Institute of Marine Biology, University of Hawaiʻi at Mānoa, Kāneʻohe, Hawaiʻi 96744, USA; Department of Anatomy and Cell Biology, The University of Kansas School of Medicine, Kansas City, Kansas 66160, USA

**Keywords:** Anthozoa, *Montipora*, *Nematostella*, body plan evolution, Hox, segmentation

## Abstract

Segmentation of the gastric cavity is a synapomorphic trait of cnidarians of the class Anthozoa (corals and sea anemones), with different clades forming distinct numbers of segments. In the starlet sea anemone *Nematostella vectensis*, for example, eight bilaterally positioned gastric segments are generated by the action of a group of Hox-Gbx genes in the developing larval endo-mesoderm. Still, given the range of segment numbers observed in different anthozoans, it remains unclear whether this Hox-Gbx module is evolutionarily conserved and how it might be deployed to generate different numbers of segments. Here, we systematically interrogate the role of Hox-Gbx genes during development of the rice coral *Montipora capitata*. We first characterize the temporal sequence of segmentation in *M. capitata* juveniles and then combine transcriptomic profiling and *in situ* hybridization to identify three conserved homeobox-containing genes, *McAnthox8*, *McAnthox6a.1* and *McGbx*, which are collectively expressed in the developing endo-mesoderm prior to and during segment formation. The expression boundaries of these genes prefigure the positions of the first six segment boundaries, similar to their *Nematostella* homologs. Further, we show that chemical inhibition of BMP activity at the planula stage abolishes the expression of Hox-Gbx genes, leading to the formation of an unsegmented gastric cavity. These findings demonstrate the existence of a functionally conserved Hox-Gbx module in evolutionarily divergent anthozoan species, suggesting that the last common ancestor of all anthozoans likely utilized a similar genetic toolkit to axially pattern the endo-mesoderm into metameric subunits.

## Introduction

Cnidarians are traditionally considered to be morphologically simple, radially symmetric organisms. However, adult cnidarians of the class Anthozoa, including corals and sea anemones, display bilateral symmetry in their arrangement of gastrodermal folds known as mesenteries (*1, 2*). These epithelial structures bridge between the body column and the pharynx, serve as the primary location for nutrient uptake and gametogenesis, and subdivide the gastric cavity into a series of interconnected segments (*3–7*). The formation of gastric segments is a prominent trait in modern corals and sea anemones as well as their extinct Cambrian ancestors such as Rugosa and Tabulata, suggesting that similar features existed in the last common ancestor of all anthozoans (*8–12*).

The molecular and cellular mechanisms that drive endo-mesodermal segmentation have been interrogated in a single anthozoan species, the starlet sea anemone *Nematostella vectensis* (*13–17*). During gastrulation, BMP signaling is asymmetrically activated on one side of the future oral pole, inducing the polarized expression of downstream homeobox-containing transcription factors (*Anthox1a*, *Anthox8*, *Anthox6a* and *Gbx*) in discrete domains of the larval endo-mesoderm (*18*). These domains collectively form a staggered pattern, likely due to distinct thresholds of gene activation downstream of BMP activity (*19*), and thereby establish an axial code that instructs the formation of four pairs of segment boundaries at their respective expression boundaries (*16*). It is worth noting that the cnidarian directive axis is not homologous to the bilaterian dorsal-ventral axis (*13, 20, 21*). The terms “dorsal” and “ventral” were historically designated as the two poles of the directive axis but do not hold comparative meaning in this context (*1*). Although BMP signaling has been implied to play a conserved role in setting up the directive axis in other anthozoans such as *Acropora*, the expression and function of downstream Hox genes have not been systematically investigated other than in *Nematostella* (*22–24*). Considering the diversity of segment numbers observed across anthozoan species, it remains unclear whether and how a functionally conserved Hox-Gbx module might regulate the bilateral segmentation process.

The rice coral, *Montipora capitata*, is a cosmopolitan species common in Hawaiʻi, where it is a primary contributor to the structure and function of local reef ecosystems (*25–27*). Despite diverging approximately 500 million years ago, stony corals (Scleractinia) and sea anemones (Actiniaria) exhibit comparable life histories and embryonic processes and likely employ similar developmental-genetic toolkits (*23, 28–30*). Consequently, *M. capitata* represents an attractive model for comparative analyses to reconstruct the ancestral function of Hox-Gbx genes in anthozoans and by extension, the cnidarian-bilaterian common ancestor. Leveraging this system, here we analyze the process of endo-mesodermal segmentation and the associated transcriptomic changes during *M. capitata* development. In addition, we combine genomics, spatial gene expression profiling, and chemical perturbation experiments to systematically interrogate the molecular functions of Hox-Gbx genes for the first time in a reef-building coral.

## Results

Despite a rich literature regarding the ecology and physiology of *M. capitata*, its early development, particularly endo-mesodermal segmentation, has remained uncharacterized. To address this, we collected *M. capitata* gamete bundles during annual spawning events at the Hawaiʻi Institute of Marine Biology (HIMB). Around summer new moons, adult *M.capitata* release buoyant gamete bundles containing sperm and eggs carrying maternally transmitted *Symbiodiniaceae* photosymbionts during mass spawning events. These gamete bundles break apart at the surface and cross-fertilize with those from other colonies (*31, 32*). By mixing gametes from multiple colonies, we observed that fertilized eggs progressed through multiple rounds of synchronized early cleavages and developed into the characteristic anthozoan “prawn-chip” blastula stage by 14hpf (**Fig. 1Ba&Ca**). Gastrulation occurred within the next 24 hours, via a combination of cell delamination and invagination, as is observed in several other corals with vertical transmission of symbiotic algae (**Fig. 1Ca**) (*33–36*). By 38hpf, embryos developed into fully ciliated oval-shaped gastrulae with clear oral openings and initiated swimming behavior (**Fig. 1Bb&Cb**). By 62hpf, teardrop-shaped planula larvae formed and continued to elongate along the oral-aboral (OA) axis, eventually achieving a rod-like morphology at 110hpf (**Fig. 1Bc&Cc, Bd&Cd**). Notably, unlike *Nematostella*, *M.capitata* planula larvae remained un-segmented, displaying no morphological divisions or cell clustering within the endo-mesoderm (**Fig. S1A-C**). We thus hypothesized that in *M.capitata,* segments only form after larval settlement.

In nature, *M.capitata* larvae settlement is triggered by multiple environmental factors, including sedimentation, light intensity, temperature, and salinity (*37–39*). Consequently, it is difficult to obtain stage-matched settlers for morphological analysis with a desired temporal resolution. To overcome this challenge, we developed an artificial settlement induction assay. The synthetic GLWamide neuropeptide from Hydra (Hym-248) has previously been shown to induce metamorphosis in several acroporid coral species with varying efficiencies, making it a promising candidate (*40–42*). When applied to 6dpf planulae at a concentration of 20µM, Hym-248 significantly altered *M.capitata* larval swimming behavior within 1 hour of treatment, shifting them from motile planulae into a sessile state (**Fig. S1D-F**, **Supplemental video 1&2**). Within 24 hours, 62.5% of larvae underwent a drastic morphological change by attaching their aboral end to the surface of the container, contracting along their oral-aboral axis, and flattening out into juvenile polyps with distinctive endo-mesodermal segments (**Fig. 1Bd-1Be**, **Fig. S1D**).

The chemically-induced settlement method described above yielded large numbers of developmentally synchronized settlers (also known as spat), enabling the characterization of the temporal sequence of segment formation (**Fig. 1D**). During early stages of settlement, we observed that the endo-mesoderm was first bisected along the directive axis into two polar segments that we named S1 and S2, with the latter being subsequently trisected into a larger polar segment S2a and two smaller lateral segments named S2b (**Fig. 1Da**). In a similar fashion, S1 was then trisected into a larger polar segment S1a and two smaller lateral segments S1b (**Fig. 1Db&Dc**). Within 24 hours of settlement induction, a total of six segments could be clearly identified, each separated by bilaterally positioned grooves corresponding to the positions of mesenteries (**Fig. 1Be**).

**Figure 1.**
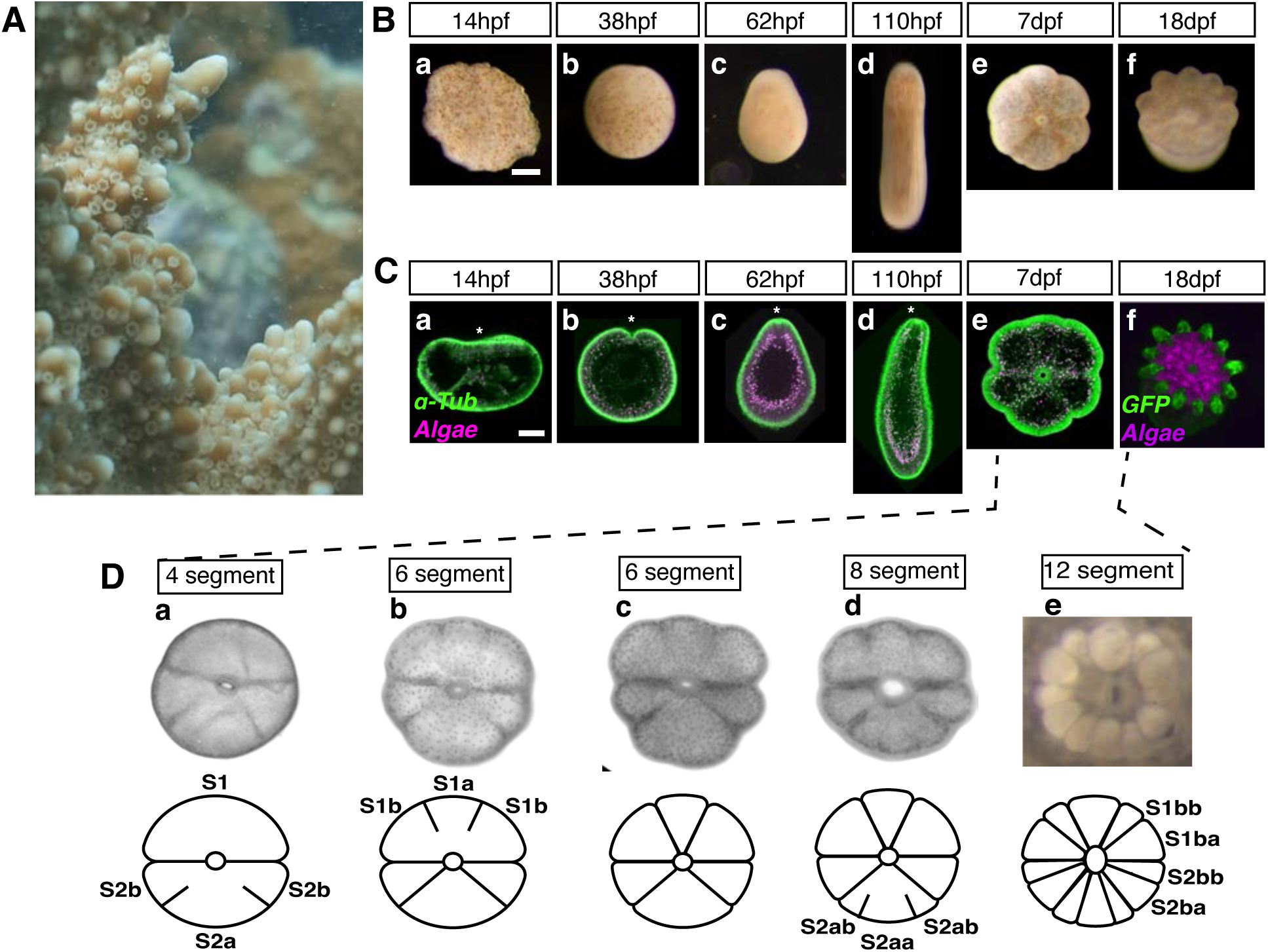
Development of the rice coral *Montipora capitata*. (A) *Montipora capitata* colony. (Ba to Bf) Brightfield images of *M.capitata* embryos collected at different developmental stages. Scale bar, 1mm. (Ca to Cf) Confocal images of *M.capitata* embryos at different developmental stages. Green, alpha-tubulin; Magenta, algal autofluorescence. Scale bar, 800 μm. Asterisks indicate the oral end. (Da to De) Oral view of segment formation during polyp development. Top panel, bright field images; bottom panel, cartoon diagrams. Segment IDs are assigned based on their order of emergence and size.

Next, a second round of segmentation began in conjunction with the elevation of the body column from the base of the spat. At this stage, polar segment S2a was further trisected into a larger central segment S2aa and two smaller lateral segments S2ab, resulting in a transient 8-segment stage (**Fig. 1Dd**). This was followed by the subdivision of S1b and S2b into the larger S1ba/S2ba and the smaller S1bb/S2bb, resulting in the formation of 12 segments (**Fig. 1De**). Unlike the first round of segmentation which occurred within 24 hours, the second round of segmentation took up to a week to complete. At approximately 10 days post settlement, we observed fully developed polyps, each possessing 12 tentacles in correspondence with the total number of segments (**Fig. 1Bf**). Based on these results, we conclude that *M.capitata* segmentation diverges from that observed in *Nematostella* in three major ways: first, there is a heterochronic shift in segmentation initiation to a later stage of development; second, the process results in the production of 12 rather than 8 segments; and third, there are discrepancies in the temporal order of segment formation (*16*).

Given these differences, we next investigated whether segmentation in *M.capitata* is driven by the same Hox-Gbx molecular code that drives the process in *Nematostella*. The available *M.capitata* transcriptome remains unannotated with a total of 63,229 protein-coding transcript models, many of which are only supported by *in silico* predictions (*43*). We therefore employed reciprocal protein blast to annotate high-confidence Hox/ParaHox candidates based on sequence homology to their *Nematostella* counterparts (**Table S1**). Despite a greater than three-fold difference in the total number of predicted protein-coding genes, the two species displayed a remarkable level of macrosynteny in terms of Hox/ParaHox organization (**Fig. 2A**). For the most part, *M.capitata* possessed a one-to-one correspondence of Hox/ParaHox genes to *Nematostella* and retained similar linkage relationships between these genes. For instance, the *Hbn*-*Rx*-*Otp* cluster on *Nematostella* chromosome 8 was found on *M.capitata* scaffold 62, with the same gene order and identical transcript orientation. The same was observed for ParaHox clusters *Mnx*-*Rough* and *B-H1*-*Ceh1*-*Hmx2*. Furthermore, each *M.capitata* Hox/ParaHox gene and the majority of its non-Hox neighbor genes could be confidently mapped to the same *Nematostella* chromosome, suggesting extensive conservation of gene arrangement at the chromosome level despite a dramatic expansion of gene number in *M.capitata* (**Table S2**).

**Figure 2.**
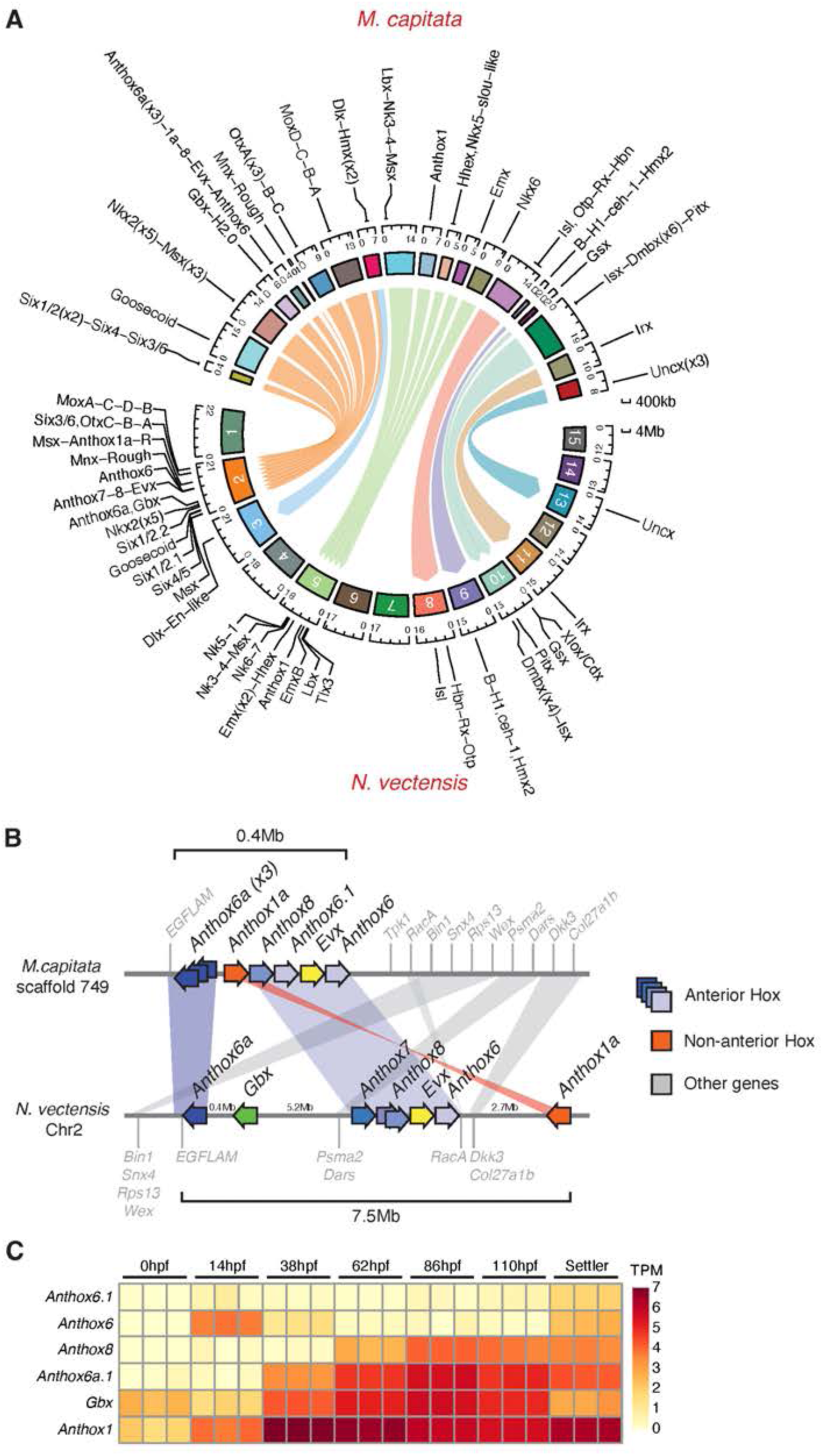
The genomic arrangement and temporal expression patterns of *M.capitata* Hox-Gbx genes. (A) Macrosynteny of homeobox containing genes between *Nematostella* and *M.capitata*. *M.capitata* scaffolds and their corresponding *Nematostella* chromosomes are connected via colored ribbons. (B) Genomic arrangement of the Hox cluster between the two species. (C) Hox-Gbx expression during *M.capitata* developmental time course.

Using this approach, we also identified a highly compacted *M.capitata* Hox cluster, consisting of three *Anthox6a* paralogs (named *McAnthox6a.1*, *6a.2* and *6a.3*), *McAnthox1a*, *McAnthox8*, two *Anthox6* paralogs (named *McAnthox6.1* and *McAnthox6.2*) and *McEvx*, located within a 0.4Mb genomic region (**Fig. 2B**). All genes shared the same transcriptional direction except the *Anthox6a* paralogs. Furthermore, this compact Hox cluster was conserved across multiple scleractinian species, in both robust and complex corals, whereas a more dispersed Hox cluster was prevalent in actiniarian species such as *Nematostella vectensis* and *Scolanthus callimorphus*, possibly due to lineage-specific chromosomal rearrangements (**Fig. S2**). Interestingly, *McGbx* was located on a different scaffold than the Hox cluster. Although 75% of all protein-coding genes on the *McGbx* scaffold (scaffold 362) and 95% of all protein-coding genes on the *McHox* scaffold (scaffold 749) possessed homologs on the same *Nematostella* chromosome (Chr2), our current data is insufficient to support linkage between *McGbx* and the *McHox* cluster in *M.capitata* (**Fig. 2B**, **Table S2**). Nonetheless, the existence of a highly organized *M.capitata* Hox cluster raises the possibility that many of these genes are coordinatively regulated and function during the same biological process.

To investigate the expression and biological function of Hox/Gbx genes in *M. capitata*, we next performed transcriptome profiling across seven developmental time points (0hpf, 14hpf, 38hpf, 62hpf, 86hpf, 110hpf and the settler stage). Intriguingly, we found that Hox/Gbx genes were expressed in a distinctive temporal sequence which preceded the formation of morphological segments in settled juveniles (**Fig. 2C**). For instance, *McGbx* mRNA was maternally deposited. *McAnthox6a.1* initiated expression at the gastrula stage (38hpf), followed by *McAnthox8*, who first became detectable in planula larvae (62hpf). The expression of all three genes persisted until settlement. Two *Anthox6* paralogs, *McAnthox6* and *McAnthox6.1*, were specifically expressed at the time of settlement. The other genes within the Hox cluster, including *McAnthox6a.2*, *McAnthox6a.3* and *McEvx*, were undetectable across all time points.

Taken together, the observations above suggest that each *M. capitata* Hox gene is subject to distinct temporal control, despite their close physical proximity in the genome. To examine their spatial expression patterns, we next performed *in* situ hybridization (ISH) using digoxigenin-labeled full-length RNA probes across different developmental stages (**Fig. 3**). Interestingly, *McAnthox8*, *McAnthox6a.1* and *McGbx* exhibited spatially restricted expression patterns within the unsegmented endo-mesoderm of planula larvae. From transverse sections at 110hpf, we found that *McAnthox8* occupied around 25% of the endo-mesoderm, *McAnthox6a.1* occupied around 50% of the endo-mesoderm and *McGbx* occupied more 75% of the endo-mesoderm (**Fig. 3F, L, R**). Collectively, these genes were expressed in a staggered fashion, with their expression domains prefiguring the position of the first six segment boundaries to form post-settlement. These observations indicate that the molecular identities of segments are established prior to the formation of physical segment boundaries, reminiscent of the Hox-Gbx dependent segmentation process in *Nematostella*.

**Figure 3.**
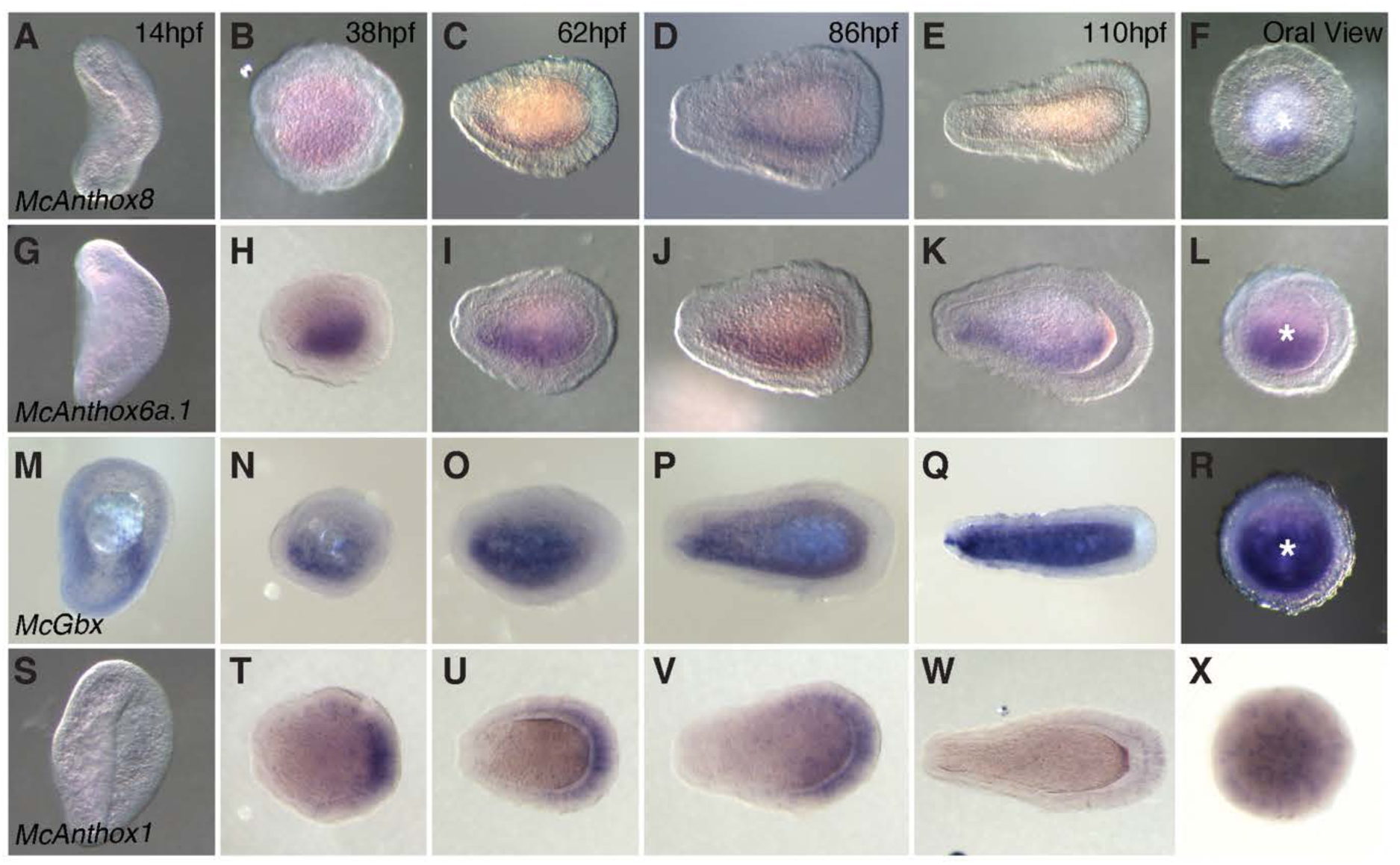
The spatial expression patterns of *M.capitata* Hox-Gbx genes. (A-F) Expression of *McAnthox8*. (G-L) Expression of *McAnthox6a.1*. (M-R) Expression of *McGbx*. (S-X) Expression of *McAnthox1*.

To determine the function of the Hox-Gbx network in *M.capitata* segmentation, we next employed chemical perturbations to disrupt its upstream regulation. The expression of *Nematostella* Hox/Gbx genes have been shown to be BMP-dependent, as they are the direct targets of Smad1/5/8 (*44*). Given that BMP components display comparable expression patterns between *Nematostella* and various acroporids, we hypothesized that inhibition of the BMP pathway in corals should also disrupt Hox/Gbx expression (*22, 23*). To inhibit BMP signal reception, we treated settlement-competent *M.capitata* larvae with either Dorsomorphin or LDN-193189, compounds which specifically target BMP type I receptors (**Fig. 4B**) (*45–47*). In clear contrast to DMSO-treated controls, Dorsomorphin-treated larvae failed to undergo segmentation upon induction of settlement, leading to the formation of spats with a hallow gastric cavity lacking mesenteries (**Fig. 4C, E, F**, **Video S3&S4**). Surprisingly, LDN-193198 treated larvae failed to respond to settlement induction with Hym-248, and thus no settlers were found 24 hours post induction (**Fig. 4D**). This may be due to the higher potency of LDN-193198, which could lead to pleiotropic effects.

**Figure 4.**
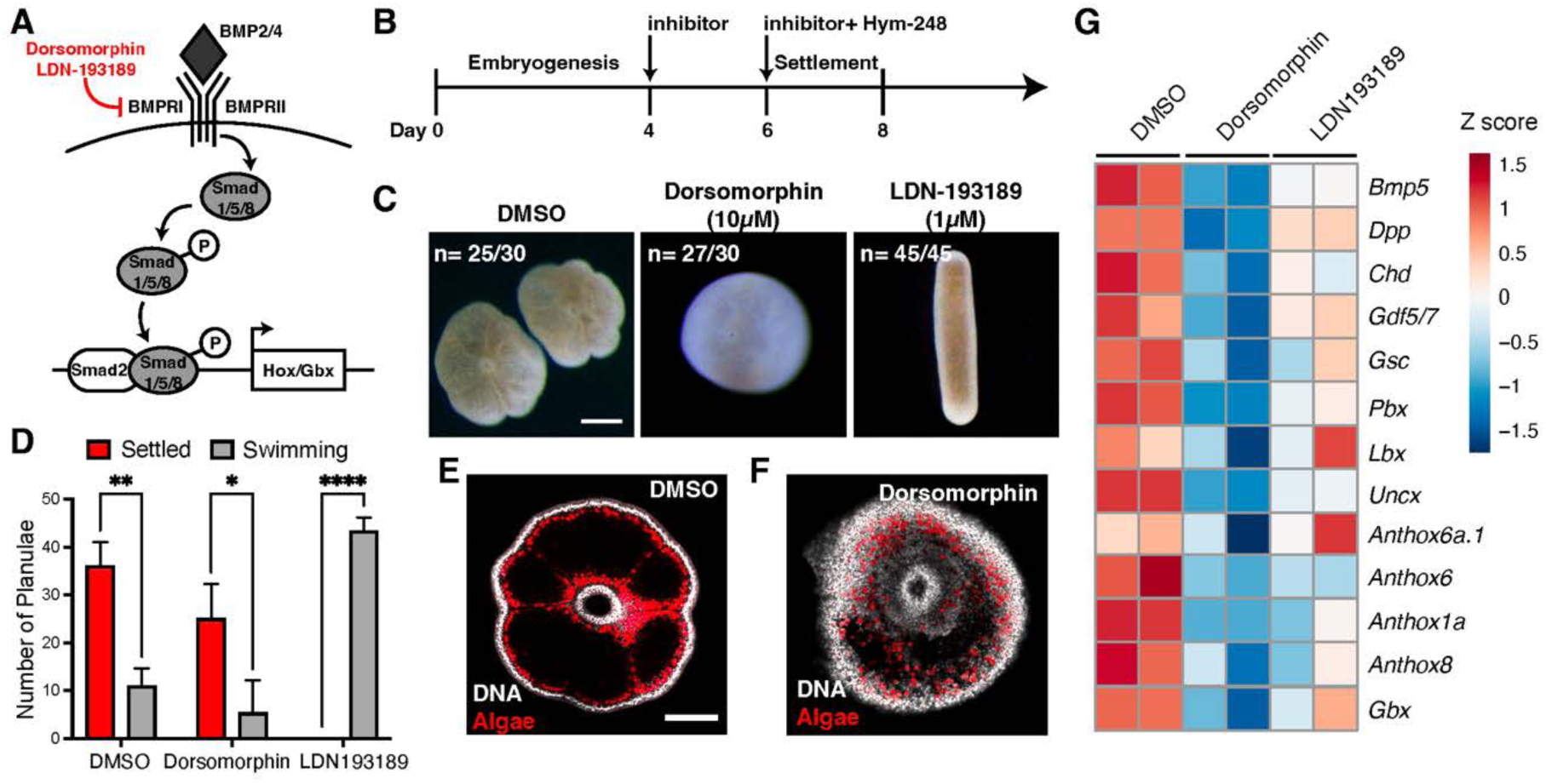
Chemical inhibition of the BMP signaling pathway results in segmentation defects in *Montipora capitata*. (A) Diagram illustrating the key components of the canonical BMP signaling pathway. Chemical inhibitors Dorsomorphin and LDN-193189 specifically target type I BMP receptor and block the activation of downstream target genes including the Hox-Gbx complex. (B) Diagram of inhibitor treatment. Both inhibitors were administered on to 4dpf larvae for 2 days prior to settlement induction using Hym-248. The morphology of settled spats as well as the percentage of settlement was documented on day 8. (C) Representative brightfield images of coral spats after DMSO, Dorsomorphin and LDN-193189 treatment. Scale bar, 1mm. (D) Percentage of larvae that settled under different treatment conditions. (E) Oral view of DMSO treated settler. (F) Oral view of Dorsomorphin treated settler. Scale bar, 100μm. (G) Expression heatmap of BMP components and their downstream targets in DMSO, Dorsomorphin and LDN-193189 treated larvae 2 days after settlement induction.

To validate the results above, we next examined the expression of BMP pathway components as well as Hox/Gbx genes in Dorsomorphin-treated larvae. Compared to controls, Dorsomorphin treatment uniformly suppressed the expression of *Bmp5*, *Dpp*, *Chordin* and *Gdf5-like*, and abolished the expression of key Hox/Gbx genes including *McAnthox6a.1*, *McAnthox8* and *McGbx* (**Fig. 4G**). These observations were further confirmed using *in situ* hybridization (**Fig. S3**). Together, our results indicate that *M.capitata* Hox/Gbx genes are expressed in patterns that correspond to presumptive segment boundaries and that the process of morphological segmentation is dependent on target gene activation downstream of BMP signaling.

Segment number and tentacle arrangement have long been utilized as important features to guide the classification of anthozoans (*48, 49*). Both stony corals and sea anemones belong to the subclass Hexacorallia, and typically possess hexameric numbers of segments (*50, 51*). As an exception, burrowing sea anemones of the family *Edwardsiidae*, including the popular genetic model *Nematostella*, possess eight complete mesenteries and display octameric symmetry, implying these animals are relatively derived. Paradoxically, molecular phylogenies based on ribosomal DNA suggests that the *Edwardsiidae* should be deeply nested within the major branch of Actiniaria (*52*). The parallels between segmentation in *M.capitata* and *Nematostella* prompted us to revisit the evolution of segmental body plans in Hexacorallia. We propose that the differences in segment number reflect a heterochronic shift during development (**Fig. 5**). On one hand, *M.capitata* settlers undergo a transient *Nematostella*-like 8-segment stage during the second round of segmentation (**Fig. 1Dd**). On the other hand, *Nematostella* polyps will eventually develop four additional mesenteries near the oral disk at around 60dpf. However, none of these fully subdivides the original segments and thus these are classified as incomplete mesenteries (**Fig. S4**). As such, a second round of segmentation happens much later in *Nematostella*, resulting in an incomplete 12-segment pattern. Most importantly, the core molecular logic for these divergent segmentation processes appears largely identical, with different Hox-Gbx genes establishing molecular territories prior to the emergence of physical boundaries. This deeply conserved Hox-Gbx code was likely already present in the last Hexacorallian common ancestor and could be readily modified to generate diverse segment patterns in corals and sea anemones.

**Figure 5.**
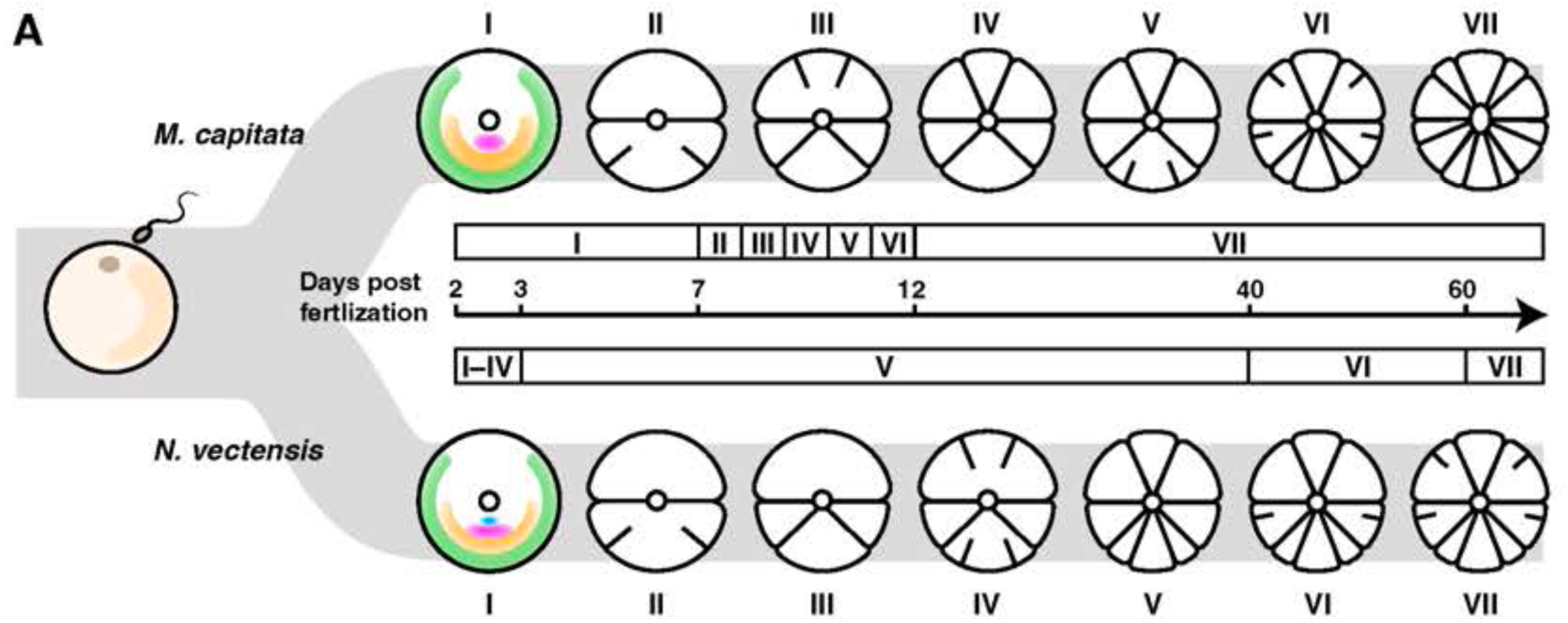
Heterochrony of segmentation sequence between *Montipora capitata* and *Nematostella vectensis*. (A) Schematics demonstrating the sequence of segment formation between the two Anthozoan species.

## Discussion

Bilaterian Hox genes often exist in close proximity to one another along the chromosome, forming what are widely known as Hox gene clusters (*53, 54*). Remarkably, the order of Hox genes along the chromosome often reflects the temporal and spatial order of gene expression during anterior-posterior axis patterning, a phenomenon called collinearity (*54–56*). In the rice coral *Montipora capitata*, we identified a highly organized Hox cluster, consisting of seven Hox genes and the homeobox-containing gene *Evx*, all located in tandem within a 0.4Mb genomic region. Comparing to other sequenced anthozoans, the *McHox* cluster possesses the largest number of genes with the smallest average intergenic distance (**Fig. S2**) (*57–63*). This is partially due to a lineage-specific duplication of the Hox gene *Anthox6a*. Three *Anthox6a* paralogs were identified in *M.capitata*, of which *McAnthox6a.2* and *McAnthox6a.3* were undetectable across all developmental stages despite their physical proximity to the highly expressed *McAnthox6a.1* (**Fig. 2B&C**). As a result, the biological significance of this gene duplication event remains unknown. We cannot exclude the possibility that *McAnthox6a.2* and *McAnthox6a.3* are pseudogenes that have lost their promoter activity, similar to the pseudogene *Anthox9* in *Nematostella* (*17*). Alternatively, these genes could be in the process of neofunctionalization and participate in different biological processes other than segmentation.

The genomic architecture of Hox genes is intimately linked to their functions during body axis establishment (*64–67*). It is postulated that the tandem localization of bilaterian Hox genes is evolutionarily favorable as it minimizes the regulatory elements required in order to achieve coordinated spatial-temporal activation (*53, 68, 69*). For cnidarians, however, the functional significance of gene clustering remains unknown, as our analysis failed to support the existence of spatial or temporal collinearity of Hox genes in *M.capitata* or *Nematostella*. We hypothesize that the Hox clusters identified in modern anthozoans were formed by ancient gene duplication events.

The relatively small size of the anthozoan cluster may have protected it against dispersions caused by chromosome-level rearrangements and thus preserved local gene linkage. Consequently, these genes do not necessarily possess a shared regulatory architecture and are not functionally constrained to participate in the same biological process. Instead, they can undergo independent neofunctionalization by acquiring novel regulatory elements within their individual promoters. This is in line with the absence of a cnidarian CTCF homolog, a crucial component for the formation of long-range genomic interactions and topological associated domains (*61*). How temporally and spatially coordinated gene expression can be achieved in this scenario remains an interesting and open question.

The unique segmentation pattern observed in the burrowing sea anemones has attracted the attention of zoologists since the late 19^th^ century. In 1891, McMurrich analyzed the developmental sequence of segmentation in the brooding sea anemone *Aulactinia stelloides* and *Rhodactis sanctithomae* and reached the conclusion that many actiniarian larvae undergo a universal “Edwardsia” stage with eight mesenteries, before developing additional mesenteries to complete the hexamerous body plan (*70*). These observations led to two different hypotheses, with the family *Edwardsiidae* representing either an ancestral form of all anthozoans or a derived form due to neoteny (*1, 70, 71*). Considering the existence of Octocorallia (soft corals), which also display octameric symmetry, McMurrich and others proposed *Edwardsiidae* as a primitive form of Hexacorallia that reflects a transitional phase between the two types of bauplan (*1, 70*).

A distinct theory was proposed in 1966 by Cadet Hand, who considered the *Edwardsiidae* body plan to be a derived trait (*71*). Modern phylogenetic reconstruction using ribosomal DNA sequences rejected the basal position of *Edwardsiidae* and instead placed them among other actiniarians, suggesting they indeed possess a derived body plan due to heterochronic development (*72, 73*). The analysis of Hox-Gbx expression presented in this work reveals homologous segment identities between *M.capitata* and *Nematostella* and provides a molecular explanation of heterochrony between these two representative species (**Fig. 5**). In both species, *Anthox6a*, *Anthox8* and *Gbx* instruct the sequential formation of three pairs of segment boundaries bilaterally positioned along the directive axis. The key difference is the lack of *McAnthox1a* expression during the first round of segmentation in *M.capitata*. In *Nematostella*, *NvAnthox1a* is co-expressed with *Anthox6a*, *Anthox8* and *Gbx* prior to segmentation, occupying an area corresponding to the future polar segment S5. Given the transient *Nematostella*-like 8-segment stage during the second round of segmentation in *M.capitata*, it is likely that *McAnthox1a* is functioning in a similar fashion, albeit much later compared to the other three genes (**Fig. 1D**). Due to the difficulty in chemical fixation of fully settled *M.capitata* polyps, we were unable to confirm the expression pattern of *McAnthox1a* by *in situ* hybridization. However, at the transcriptomic level, *McAnthox1a* was drastically downregulated in Dorsomorphin-treated settlers, in accordance with the rest of segmentation-related Hox/Gbx genes (**Fig. 4G**). Taken together, our findings strongly suggest that the heterochronic deployment of a conserved Hox-Gbx module contributes to the divergent adult body plans observed between *Edwardsiidae* and other anthozoans.

In summary, this work establishes *Montipora capitata* as a powerful comparative system to understand the evolution of Hox genes and body segmentation in anthozoans. In comparison to *Nematostella*, *M.capitata* displays some unique developmental features such as cell delamination during gastrulation and rapid endo-mesodermal segmentation during settlement. Despite these differences, *M.capitata* possesses a conserved set of Hox-Gbx genes (**Fig. 2**). Three of the genes, *McAnthox8*, *McAnthox6a.1* and *McGbx* are coordinately expressed in the developing endo-mesoderm in a staggered fashion during the segmentation process (**Fig. 3**). Perturbation of the BMP signaling pathway reduces the expression of these Hox-Gbx genes and abolishes segment formation during metamorphosis (**Fig. 4G**). Future work should focus on the dynamics of Hox-Gbx expression in post-metamorphic *M.capitata* polyps and on developing reverse genetic tools such as shRNA knockdown and CRISPR/Cas9 mutagenesis to provide functional validation of these genes. Nonetheless, our observations in *M.capitata* suggest that the expression and regulation of Hox-Gbx genes are highly conserved within the anthozoan lineage and are intimately linked to the endo-mesodermal segmentation process. A similar genetic program may have existed in the anthozoan common ancestor to generate metameric features 500 million years ago.

## Acknowledgement

This work was done in part to fulfill S.H.’s Ph.D. requirement. We would like to thank Josh Hancock for sample collection and logistical support. E.R.H was partially supported by an Emerging Research Organisms Grant from Society for Developmental Biology (102511/SDB0003), C.D. and E.M. are supported by the Paul G. Allen Family Foundation and National Science Foundation (IOS-2041401). This work was generously supported by the Stowers Institute for Medical Research. Coral gametes were collected under permits from the Hawaiʻi Division of Aquatic Resources to HIMB/CD (SAP 2020-25, SAP 2021-33, SAP 2022-22, SAP2023-31).

## Author Contributions

S.H., E.R.H. and M.C.G designed the experiments, E.M. and C.D. collected the samples, S.H., E.R.H., E.H. and L.E. performed the experiments. S.H., E.R.H, S.C. and S.R. performed data analysis. S.H., E.R.H. and M.C.G. wrote the manuscript. All authors have read and agreed to be listed on this manuscript.

## Material and Methods

### Sample collection and animal handling

*Montipora capitata* gamete bundles were collected from Kāneʻohe Bay (Oʻahu, Hawaiʻi) during the summer spawning events between 2020 and 2023. Gamete bundles from wild colonies were collected under ambient conditions, returned to HIMB, and allowed to break up and fertilize following established best practices (*74*).

Settlement induction was performed using two independent methods. Starting from 6dpf, planula larvae were placed into settlement bins with aragonite plugs covered in crustose coralline algae in running seawater. Another batch of planula larvae were treated with 20µM Hym-248 neuropeptide (EPLPIGLW-amide, NovoPro, #304370) in 35ppt artificial seawater. Metamorphosed juveniles were kept in 26°C incubators with 12-12 lighting and daily water exchange to ensure their normal growth.

### Metamorphosis induction experiment by synthetic neuropeptide

Larvae from 3-6 dpf were assessed for the dose-response of Hym-248 neuropeptide activity; between 10-30 planula larvae were placed per well to sterile 6-well plates with 10 ml of filtrated seawater at 26°C, without additional substrate for attachment.

Hym-248 was added to final concentration of 1μM and 10μM (stock solution 2.7mM) in larvae of 3 – 6 dpf, following a previous concentration used in *Acropora palmata* (*41*). Larvae were maintained with the neuropeptide during larval competence development. At 3 dpf there was no significant effect of the neuropeptide Hym-248 in both concentrations. Once the planulae had raised searching behavior at 4 dpf, >10% of larvae settled at 10μM but not 1μM concentration. At 5 dpf, larvae became more active than in earlier stages getting a higher percentage of the settlement at 10 μM (48%) showing dose-response of Hym-248. At 6 dpf, we introduced a concentration of 20 μM in the induction experiment, due to the raising response at 5 dpf. Both concentrations (10 and 20 μM), induced more than 60% of settlement; however, 20 μM had a slightly stronger effect than 10 μM, for further experiments we used 20 μM as final concentration, without affecting the survival rate.

### BMP-inhibitors treatment

Planulae larvae of 3dpf were placed on 6 well plates with 10 ml of filtrated seawater at 26°C. The planulae larvae were maintained with 10μM of Dorsomorphin (2.5 mM stock solution, Cayman chemical, *#*11967) and 1uM of LDN193189 (10mM stock solution, Sigma-Aldrich, # SML0559) and DMSO as controls as well. Planula larvae were maintained with the treatment for 3 days before and after the induction with Hym-248. At 10 dpf settlement and metamorphosis were quantified; finally, at 11 dpf those planulae (treatments and controls) were either fixed for ISH or placed in Trizol for localization of mRNA transcripts within whole planula larvae and the RNA collection, respectively.

### RNA collection and transcriptome profiling

Samples for the developmental time series were harvested at 0, 14, 38, 62, 86 and 110 hpf with 5 replicas per time point. For each sample, 100-200 larvae were collected in a 1.7mL Eppendorf tube and spin down at 1000 rpm to allow the removal of excess seawater. 500µL of RNA*later*® (Sigma-Aldrich, R0901) was then added to each tube and kept at -20°C. Each settlement replicate was collected by pooling 80-100 settled larvae into a 1.7mL Eppendorf tube and spinning down at 1000 rpm to allow the removal of excess seawater. 500µL of TRIzol reagent was added to the tube and kept at -20°C.

RNA extraction was performed using Direct-zol™ RNA Miniprep Plus Kit (Zymo, R2071) following the manufacturer’s instruction. RNA quality was checked using Bioanalyzer (Agilent) and the three replicas with the best RNA integrity were submitted for sequencing.

### Transcriptome assembly and RNA-seq analysis

Published *M.capitata* genome, transcriptome, and proteome were downloaded from http://cyanophora.rutgers.edu/montipora (*43*). However, this version of the transcriptome only contains CDS sequences. To better quantify RNA-seq signal, we performed de novo transcriptome assembly using published *M.capitata* RNA-seq datasets of various stages. RNA-seq libraries for different timepoints were pooled and sequenced using Illumina Nextseq 500 instrument with 75bp read length. Raw reads were demultiplexed into Fastq format allowing up to one mismatch using Illumina bcl2fastq2 software v2.18. Genome and transcriptome builds were downloaded from the Shumaker et al manuscript [1] and aligned using STAR aligner v2.7.3a. During alignments, reads that aligned to Cladocopium and Durusdinium algae were removed for downstream analysis. Finally, TPM values were generated using RSEM v1.3.0. Differential gene expression (DGE) analysis were performed using R package edgeR and heatmaps were generated using R package pheatmap (*43*).

### Identification of homeobox-containing genes in *Montipora capitata*

To identify homeobox-containing genes in *Montipora capitata* genome, a customized BLASTP database was set up locally (*75*). Protein sequences of *Nematostella* homeobox-containing genes were blasted against published *M.capitata* proteome (http://cyanophora.rutgers.edu/montipora) with an E-value cutoff of 1e-20 (*43*). For each gene, the top three *M.capitata* candidates were selected and reciprocally blasted against *Nematostella* transcriptome using the SIMRbase web interface (https://genomes.stowers.org/) (*61*). The *M.capitata* gene/genes with the lowest E-value were then named according to the naming system in *Nematostella*. The gene ID, homology, and E-value of *M.capitata* homeobox-containing genes are listed in Table S1. The gene ID, genomic location, homology, and E-value of all *M.capitata* genes used for synteny analysis were listed in Table S2.

### Immunohistochemistry and <u>W</u>holemount in <u>s</u>itu <u>H</u>ybridization (WISH)

Samples for the developmental time series were harvested at 0, 14, 38, 62, 86 and 110 hpf in 15mL falcon tubes. Majority of the sea water was removed before adding 12mL 4% Paraformaldehyde diluted in seawater and fixed at room temperature for 1.5 hours on a shaker. Animals were then washed 4 times in 1x PBST and dehydrated through methanol gradients. Samples were stored in 100% methanol at -20°C. Prior to immunostaining or ISH, animals were first washed for 10 mins in 100% xylene to remove the lipid inside the tissue. The sample was then washed back into 100% methanol twice and rehydrated through the methanol/PBST gradient. The following antibodies and dyes were used for immunostaining: mouse anti-α-tubulin (Sigma, 1:1000), SiR-DNA (Cytoskeleton, 1:1000), Goat-anti-mouse secondary 647 (Life Tech, 1:1000). For WISH, animals were first treated with 3% H2O2/methanol at room temperature for 1 hour, then digested using 80µg/mL Protease K at room temperature for 2 mins and quickly transferred into 4% Paraformaldehyde in 1x PBST for 1 hour. Hybridization was carried out with 1ng/μL digoxigenin (DIG)-labeled probes at 60°C for up to 60 hours. Probe synthesis was carried out following published protocols (*76*). Samples were washed through regular hybridization buffer/SSC series and incubated in 1:1000 pre-absorbed anti-DIG-AP fab fragment (Roche)/5% sheep serum/1% blocking reagent (Roche)/1% DMSO/TNT at 4°C overnight. Colorimetric reaction was performed using the NBT/BCIP method at room temperature overnight. Samples with sufficient signal were fixed and quickly rinsed in 66% ethanol to remove background and mounted in Scale A2 (2M Urea, 75% Glycerol) buffer before imaging. All primers used for probe synthesis are listed in Table S3.

### Imaging and image processing

Immunostaining samples were imaged using Zeiss 700 upright confocal microscope with ZEN software. Colorimetric WISH samples and bright filed images of *M.capitata* polyps were imaged under Leica M165 FC stereo microscope and Axiovert 206 MOT. Image processing was performed using Fiji following standard procedures. Unprocessed raw images can be accessed from the Stowers Institute for Medical Research Original Data Repository.

## Data availability

All NGS data has been deposited to the Gene Expression Omnibus (GEO) and can be accessed at: (https://www.ncbi.nlm.nih.gov/geo/query/acc.cgi?acc=GSE267847). The original data can be accessed from Stowers Institute for Medical Research Original Data Repository (https://www.stowers.org/research/publications/libpb-1642).

