## Supplemental figures for "An evolutionarily conserved Hox-Gbx segmentation code in the rice coral *Montipora capitata*"

[2]Hawai‘i Institute of Marine Biology, University of Hawai‘i at Mānoa, Kāne‘ohe, Hawai‘i 96744, USA

**SUPPLEMENTARY FIGURES**

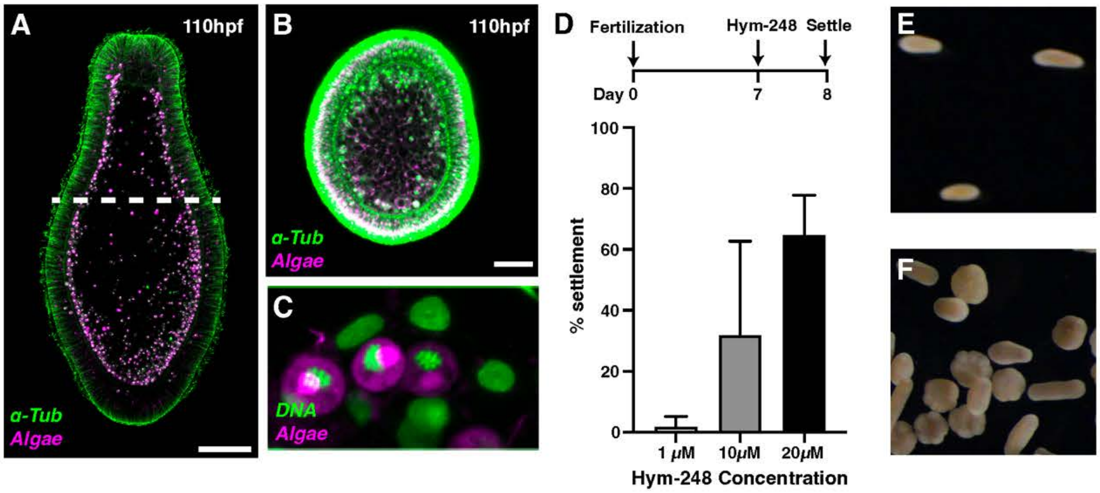


**Figure S1. Chemically induced endo-mesodermal segmentation in *M.capitata*.** (A) A sagittal section of a *M.capitata* planula at 110hpf. Dashed line indicates the relative position of transverse section in B. Green, α-Tubulin; magenta, autofluorescence from the symbiotic algae; Scale bar, 100μm. (B) A transverse section demonstrating the unsegmented endo-mesoderm at 110hpf. Green, α-Tubulin; magenta, autofluorescence from the symbiotic algae; Scale bar, 100μm. (C) Zoomed in view of the symbiotic algae inside *M.capitata* cells. Green, SiR-DNA; magenta, autofluorescence from the symbiotic algae. (D) Schematics of the settlement induction assay and quantifications of the settlement percentage across different Hym-248 concentrations. (E) Planulae morphology in DMSO control. (F) Planulae morphology in Hym-248 treated dish.

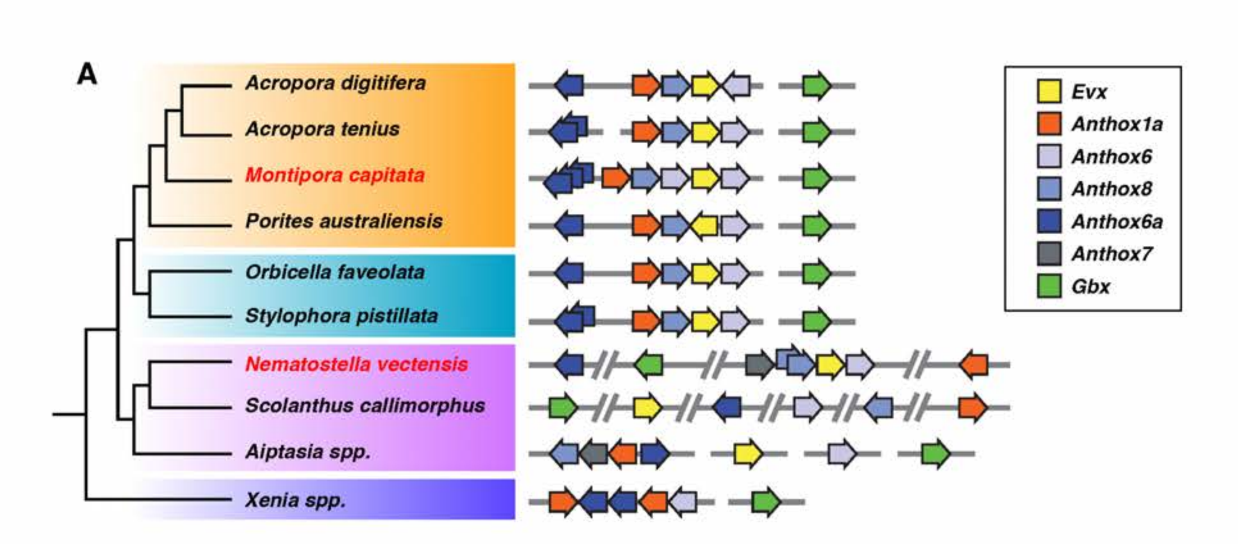


**Figure S2. Genomic conservation of the anthozoan Hox cluster.** (A) Cartoon illustrating the genetic composition and order of Hox/Gbx genes within the cluster. Each gene is color coded to match their *Nematostella* homologs. Direction of the arrow indicates the direction of transcription.

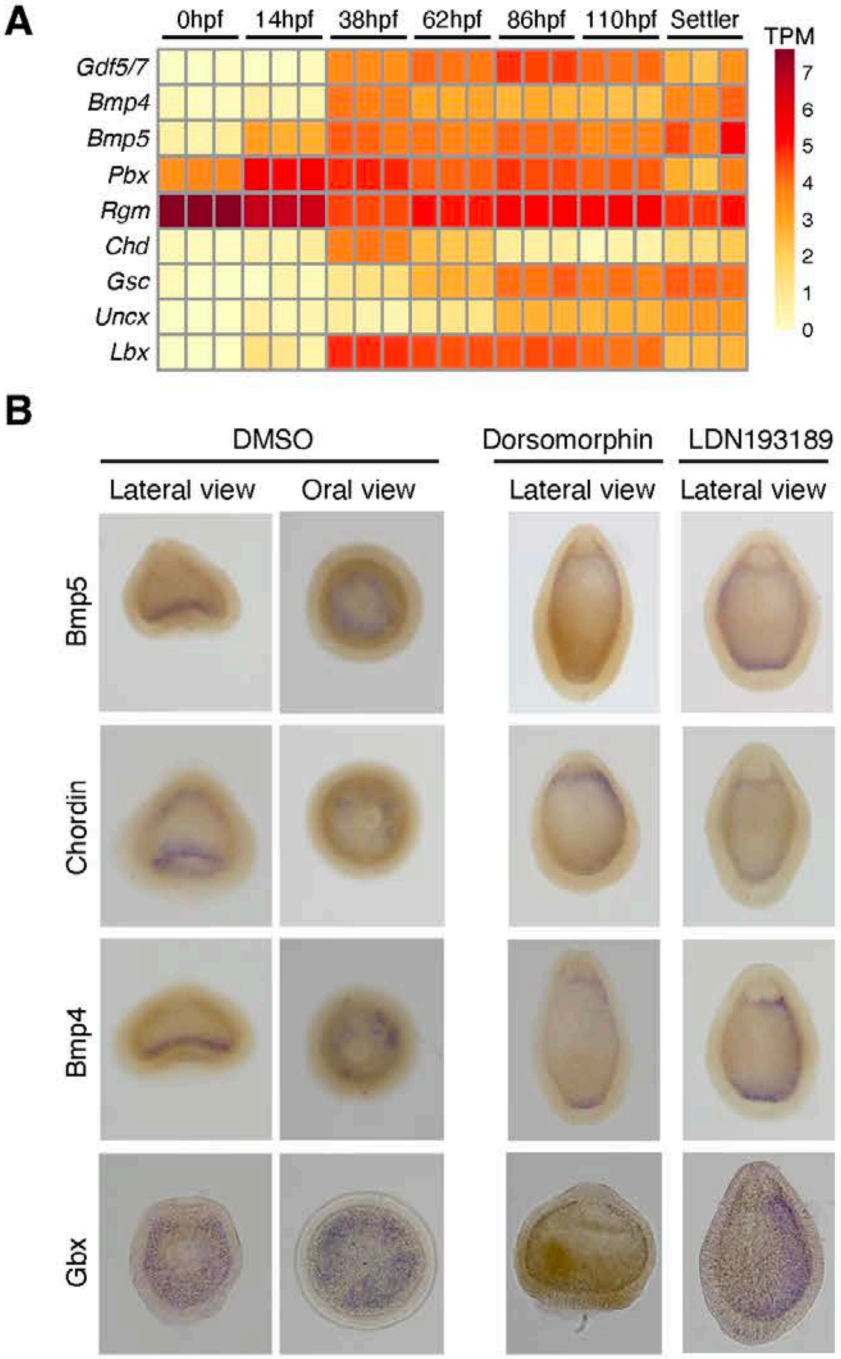


**Figure S3. Expression of BMP components and BMP target genes during *M.capitata* development.** (A) Expression heatmap of BMP components as well as their known targets across

*M.capitata* developmental stages. (B) Expression patterns of *McBmp5*, *McChordin*, *McDpp* and *McGbx* in wildtype and BMP inhibitor treated settlers. Scale bars, 100µm**.**

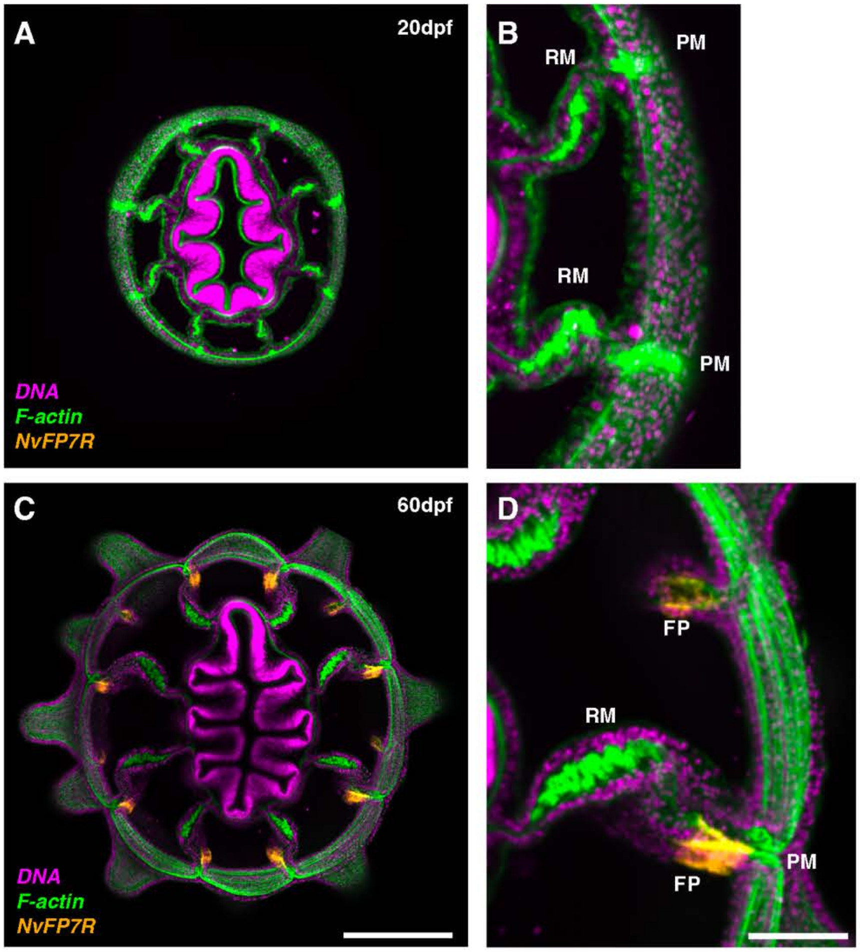


**Figure S4. The development of additional mesenteries in the starlet sea anemone *Nematostella vectensis*.** (A) Transverse view of a 20dpf *Nematostella* polyp showing the position of eight segments. Green, F-actin; magenta, DNA, orange, NvFP7R autofluorescence. (B) A zoomed in view of mesenteries at 20dpf. RM, retractor muscle; PM, parietal muscle. Green, F-actin; magenta, DNA, orange, NvFP7R autofluorescence. (C) Transverse view of a 60dpf *Nematostella* polyp showing the position of incomplete mesenteries. Green, F-actin; magenta, DNA, orange, NvFP7R autofluorescence. Scale bar, 200 µm. (D) A zoomed in view of incomplete mesenteries at 60dpf. RM, retractor muscle; PM, parietal muscle, FP, fluorescent patch. Green, F-actin; magenta, DNA, orange, NvFP7R autofluorescence. Scale bar, 100 µm.
